# Determination of Drug Sensitivity in Patient Derived Models of Breast Cancer by Multiparametric QPI

**DOI:** 10.1101/2025.10.01.674349

**Authors:** Edward R. Polanco, Tarek E. Moustafa, Ozlen Balcioglu, Sandra D. Scherer, Emilio Cortes-Sanchez, Sophie Remick, Benjamin T. Spike, Alana L. Welm, Bryan E. Welm, Philip S. Bernard, Thomas A. Zangle

## Abstract

Functional precision oncology seeks to match patients with effective therapies by empirically testing patient-derived samples for drug sensitivity in the laboratory. However, existing approaches require significant sample expansion time and expense prior to analysis, rendering them impractical for routine clinical testing. Quantitative phase imaging (QPI) provides a potential path forward by directly measuring responses at single cell resolution without the need for extensive sample expansion. In previous work, we demonstrated that multiple, independent parameters of cellular response to therapeutic agents can be derived from QPI data, an approach we call multiparametric QPI (mQPI). Here, we demonstrate application of mQPI using cells from patient derived xenograft organoid (PDxO) models, as well as cells viably frozen direct from patients. Using mQPI with breast cancer PDxO models, we uncover distinct drug responses for cells originating from different anatomic sites in the same patient and resolve cellular heterogeneity of response in a model of acquired therapeutic resistance. We also show that mQPI can detect drug responses in viably frozen primary patient samples, either direct from thaw or after a short term expansion of only 2 weeks. Overall, these data provide proof-of-principle for application of mQPI to a range of sample types, including cryopreserved material direct from patients. This underscores the clinical potential of mQPI as a time- and materials-efficient alternative to current methods in functional precision oncology.

## Introduction

Precision oncology uses genomics and comprehensive sequencing to identify molecular alterations in tumors for targeted therapy.^1,2^ While this approach has improved survival for patients with some cancer types, many variants/mutations lack evidence of clinical utility and the emergence of resistance to targeted drugs is common.^3,4^ These challenges have led to an emerging discipline of precision oncology that uses patient derived models of cancer (PDMC) to determine a functional drug response for guiding patient therapy^5–8^ For example, drug responses in human breast cancers grown in patient derived xenograft (PDX) models show high agreement to responses in the patient. However, these PDX models can require up to 6 months of tumor expansion and can then only screen a few therapies. Thus, these methods are not amenable to clinical testing due to high cost, slow turnaround time, and low throughput.

In an effort to reduce reliance on *in vivo* animal testing and generate a larger library of relevant cancer models, the PDMC field transitioned to culturing patient derived xenograft cells as organoids (patient derived xenograft organoids, PDxOs)^9^ and monitoring cell growth and drug response. For *ex vivo*. For example, breast cancer PDxO models have been shown to maintain the genetics and drug response characteristics overtime. *Ex vivo* cancer cell response/resistance to different therapies can be estimated from EC50 values calculated using CellTiter-Glo (CTG), an endpoint metabolic (ATP content) assay. Although the CTG assay reflects PDxO growth, it does not provide information about heterogeneous cellular behaviors and as an endpoint assay it does not grant insight into the temporal dynamics of response. Another technique for estimating cancer cell drug responses uses microchannel resonators to measure the change in mass over time as single cells pass through the channel.^10–12^ However, since cells flow through the device as the measurements are made, microchannel resonators less suitable for studying adherent cell types.

One emerging approach for studying the temporal dynamics of cell response to therapy is quantitative phase imaging (QPI),^13^ a quantitative microscopy technique that measures changes in the optical path length as light passes through cellular material.^14^ This ‘slowing down’ of light is proportional to cell dry mass, thus, allowing QPI to measure cell growth and death of single cells as the accumulation and loss of mass over time respectively.^15,16^ QPI is label-free, facilitating the study of live cells without disrupting cell activity.^17^ This is especially important for application to clinical samples which may have limited number and that may be sensitive to labeling approaches. Initial QPI systems were based on interferometry^18,19^ while later techniques include tomography,^20^ holography,^21^ and wave front sensing.^22^ Each of these techniques comes with varying levels of precision and accuracy^23^ as well as system specific optical complexity, references, alignment challenges and computational requirements. For example, a standard brightfield microscope can be used with computational techniques that implement the transport of intensity equation to compute phase shifts from out of focus images.^24,25^ Cross-grating wavefront microscopy (CGM)/quadriwave lateral shearing microscopy (QLSI), a wavefront sensing approach, can be performed with commercially available cameras on standard research microscopes.^26–28^ Fourier ptychography, which folds images captured under a wide range of illumination to capture phase shifts at resolution beyond the diffraction limit,^29,30^ or digital holography,^31–33^ can be performed on custom or commercial purpose-built systems. Finally, differential phase contrast (DPC)^34^, a computational microscopy approach, combines four brightfield images captured using an LED array as a light source on an otherwise standard microscope to compute phase.^34,35^ Of these, DPC, in particular, offers a robust platform that is well-suited to high-throughput functional assays.

QPI has been used to quantify growth and other cellular behaviors in a variety of settings. For example, previous work used QPI to examine biomass partitioning during cell division,^15^ cell death,^36–38^ and transport of bulk material inside a cell.^39,40^ QPI has also been used to study the mechanical properties of cells such as the shear modulus^41^ and viscoelasticity.^42,43^ The dynamics of neuron growth and branching has also been previously studied with QPI.^44–46^ QPI can also be used to monitor disease progression in terms of changes in tissue structure,^47–49^. With the aid of machine learning approaches, QPI has been used to generate synthetic H&E staining,^50^ and to track organelles.^51^

A key application of QPI involves the study cell behavior in response to perturbations such as drug response.^52–54^ The high precision of QPI has enables rapid assessment of response to drug^55^ or in bioprinted organoids.^56^ In our previous work on breast cancer cell lines, we developed several parameters that leverage unique capabilities of QPI.^53^ Since timelapse QPI measures changes in mass with single cell resolution, we were able to determine the growth rate of individual cells and heterogeneity in populations as they change under therapeutic challenge. Accordingly we were also able to establish EC_50_ values by QPI, i.e. the relative concentration at which half the maximal response to therapy occurs and these values were highly concordant with those obtained by CTG.^53^ Furthermore, QPI enables depth of response (DoR) measurements as the difference between the asymptotes along a dose response curve normalized by the baseline growth rate of the control. QPI can also leverage time-series data to study the temporal dynamics of cell behavior, such as the time required to elicit a drug response, which we describe here as time of response (ToR). Taken together, these various parameters allow QPI to provide a multiparametric tool that builds a more robust picture of how cancer cells are responding to therapy.

Here, we apply QPI as a multiparametric approach for understanding how PDxO samples respond to cancer therapies. We demonstrate that QPI can identify the difference in response between PDxO models derived from different tumor sites within the same patient. We also show that QPI can be used to study heterogeneous samples and the emergence of resistant subpopulations. We also demonstrate the application of multiparametric QPI to CD45 depleted tumor cells in direct-from-thaw patient samples, and short-term expanded patient samples, which we show greatly enriches the tumor content of the sample. Toward a practical solution for precision oncology, this approach can provide empirical drug response data for solid tumors on a patient-by-patient basis with lower time and material overhead than existing methods.

## Results

### Multiparametric QPI characterizes drug response of PDXO models

We used mQPI to assess how drug treatment affects mass and growth rate of breast cancer cells from PDXO models at single cell resolution (Fig. 1). Short term two-dimensional (2D) cultures were established from PDXO models by dissociating organoids and seeding on polyethylenimine (PEI)-coated 96 well plates (Fig. 1a, S1). PEI, a positively charged polymer used for plating neural cell cultures,^57^ was found to be a suitable substrate for PDXO breast cancer cell growth and QPI imaging (Fig. S2). The PDxO models were previously established at the Huntsman Cancer Institute at the University of Utah to create a bank of PDxOs representing important clinical sub-types including triple negative breast cancer (TNBC), metastatic cancer, and treatment resistant cancers (Table ST1).^9,58,59^ Eighteen to twenty-four hours after plating in 2D, cultures were dosed with 5 different drug treatments in 6-point dose curves, and mQPI data was collected for 48 hours (Fig. 1b-i, S1) (Fig. S3, Tables ST2, ST3).

**Figure 1.**
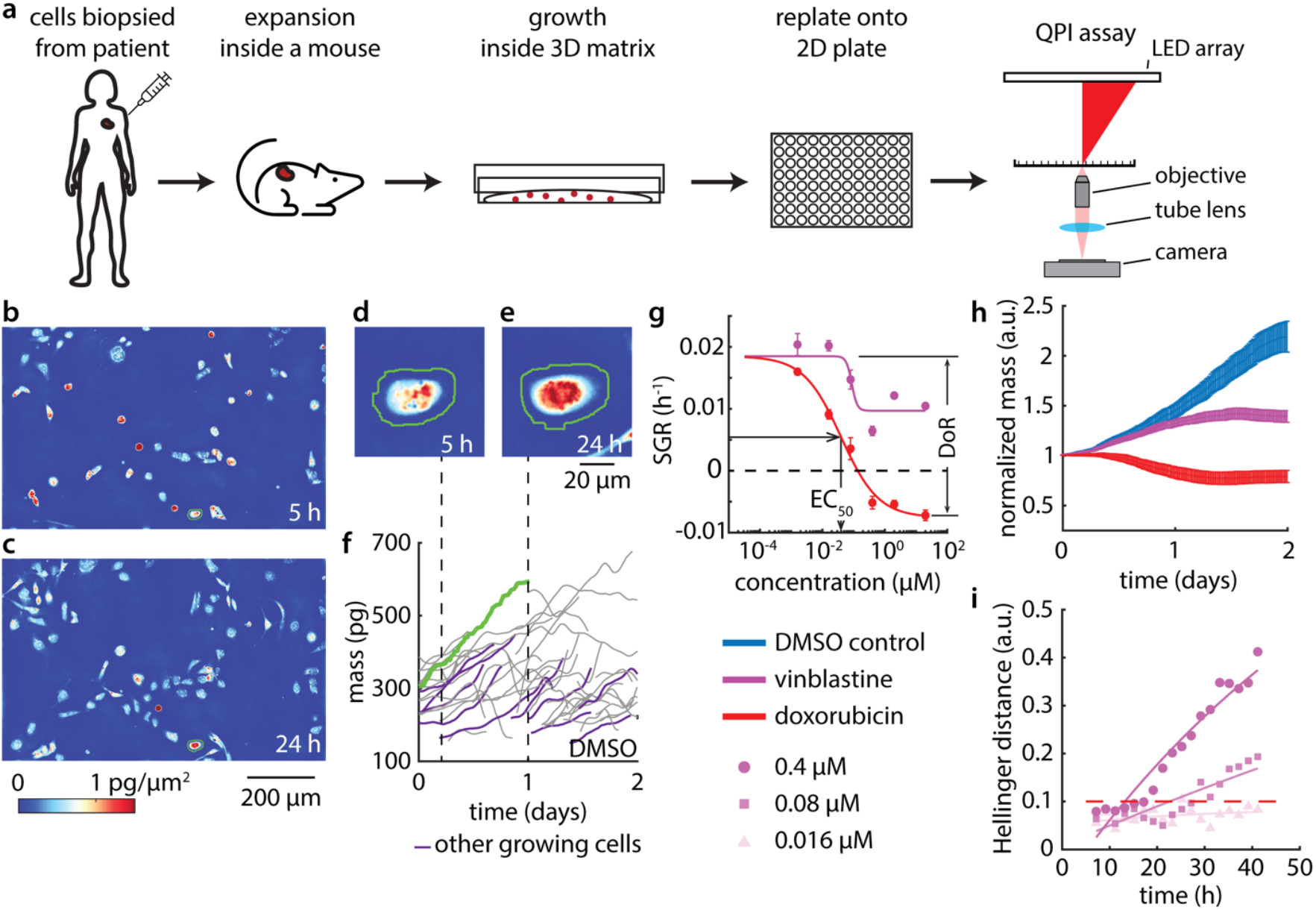
Multiparametric QPI characterizes response of PDXO model HCI-037 to cancer therapy. (a) Experiment workflow. (b-e) Images of DMSO treated HCI-037 PDXO cells in 2D culture at (b) 5 h and (c) 24 h. An individual HCI-037 PDXO cell at (d) 5 h and (e) 24 h. (f) Dry mass vs time traces for the cell in (d-e) (green line) and other proximal cells (purple). (g) Dose response plot of HCI-037 for doxorubicin and vinblastine treatments from all individual cell tracks at each drug concentration. (h) Normalized mass shows the temporal dynamics of HCI-037 response to doxorubicin and vinblastine at the EC50, showcasing a drug dependent response. (i) Hellinger distance plot for 0.4, 0.08 and 0.016 μM vinblastine.

We imaged cells for 48 h using an automatic segmentation algorithm to identify individual cells (Fig. 1b,c). The mass of each cell was determined by summing mass over the segmented area (Fig. 1d,e). Plots of mass vs. time reveal single cell growth rates (Fig. 1f), which we fit to a linear model by least-squares regression. Specific growth rate (SGR) was computed for each cell by normalizing the slope of this line by initial mass to account for the effect of cell size on growth rate.^11,53^ We then fit SGRs of each drug as a function of dose to a sigmoidal dose response curve described by a 3-parameter Hill model to extract dose response measurements, EC_50_ and depth of response (DoR) (Fig. 1g). Plots of total cell mass over time, normalized by initial mass, show the drug dependent nature of the population average temporal growth response at EC_50_. For example, some drugs, such as doxorubicin, show a large decrease in mass, starting at t = 0 h, while others, such as vinblastine, show a smaller effect on mass, with separation from control delayed to later timepoints (Fig. 1h). Each PDXO model was tested in triplicate against each drug dose. We observed low variation in control growth rate between biological replicates, indicating the repeatability of the QPI assay (Fig. S4).

Finally, to capture the time-dependence of drug effects, we compare the distribution of single-cell SGRs within each time window following treatment. For each drug-concentration pair, the difference between treatment and control was quantified using Hellinger distance^12^ and fit to a single exponential.^53^ Time of response (ToR) is then the time at which the fitted line crosses a certain threshold determined by comparison to the maximum variation observed over time in control wells (Fig. S4). ToR reveals concentration dependent temporal dynamics. For example, HCI-037 treated with increasing concentration of vinblastine show progressively faster ToR (Fig. 1i).

### Multiparametric QPI characterizes differential growth behavior between PDXO models from various sites of origin

We used multiparametric QPI to characterize the difference in response from three PDXO models derived from a single patient biopsied at different timepoints, locations, and intervening treatments. These include the primary tumor (HCI-037), bone metastasis (HCI-038), and pleural effusion (HCI-039).^9^ These different PDXO models have distinct morphologies when imaged via QPI, with HCI-037 and HCI-038 being flatter while HCI-039 is more spindle-like (Fig. 2a). Despite differences in morphology, these three PDXO models have nearly the same average SGR (Fig. 2b) and single cell growth rate distributions (Fig. S5).

**Figure 2.**
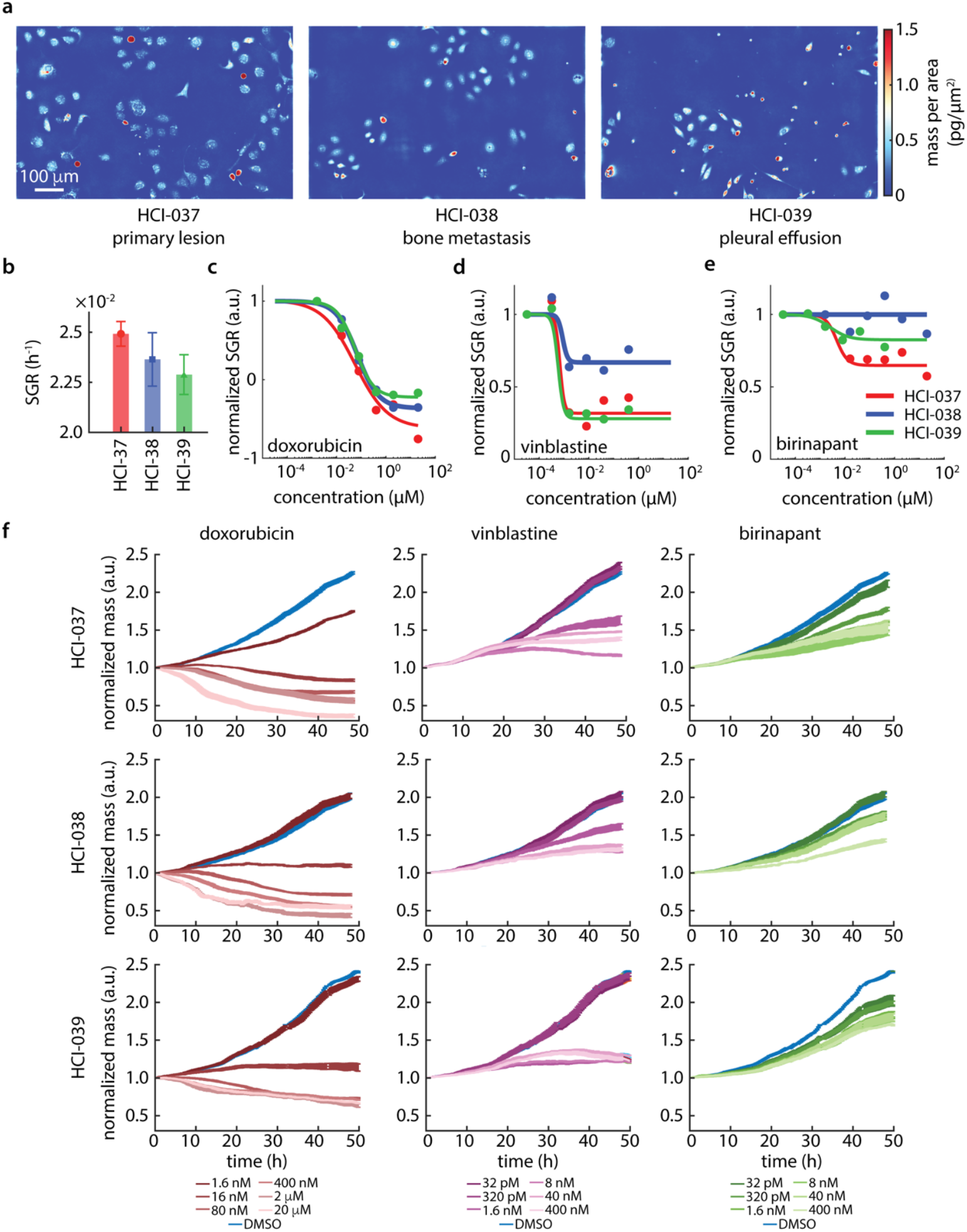
Multiparametric QPI shows differential response in tumor cells from different sites of origin. (a) Representative QPI images of three PDXO models from the same patient, HCI-037 (primary), HCI-038 (bone metastasis), and HCI-039 (pleural effusion). (b) Population average SGR for each model. (c-e) Dose response plots for each model with (c) doxorubicin, (d) vinblastine, and (e) birinapant treatment. Colors indicate model (red: HCI-037, blue: HCI-038, green: HCI-039). (f) Longitudinal measurement of mass normalized to initial masses for the same drugs as in (c-e) at varying dose (darker: higher dose). Error bars show SEM.

We measured the dose response behavior of HCI-037, HCI-038, and HCI-039 PDXO models (Fig. 2c, S5). We found that while all three PDXO models demonstrated a similar EC50 and DoR to doxorubicin (Fig. 2c), the models exhibit differential DoR in response to vinblastine (Fig. 2d) and birinapant (Fig. 2e), with the bone metastasis exhibiting a generally weaker DoR overall. This is evident in normalized mass data as well, with full response to birinapant resulting in residual growth at the end of the 48 h QPI assay (Figs. 2f, S6, S7).

### Multiparametric QPI characterizes heterogeneous growth behavior in PDXO model after selection for resistance to birinapant

We next tested two PDXO models that originate from the same tumor sample, with one cultured under standard culture conditions, HCI-027BS, and one cultured under selective pressure to enrich a population resistant to birinapant, HCI-027BR (Fig. 3a). As previously reported, these models show differential sensitivity to birinapant as PDXO.^9,59^ When grown in 2D culture for QPI, HCI-027BR grew slightly faster than the HCI-027BS (Fig. 3b). HCI-027BS also exhibited a notably more bimodal distribution than HCI-027BR (Fig. 3b).

**Figure 3.**
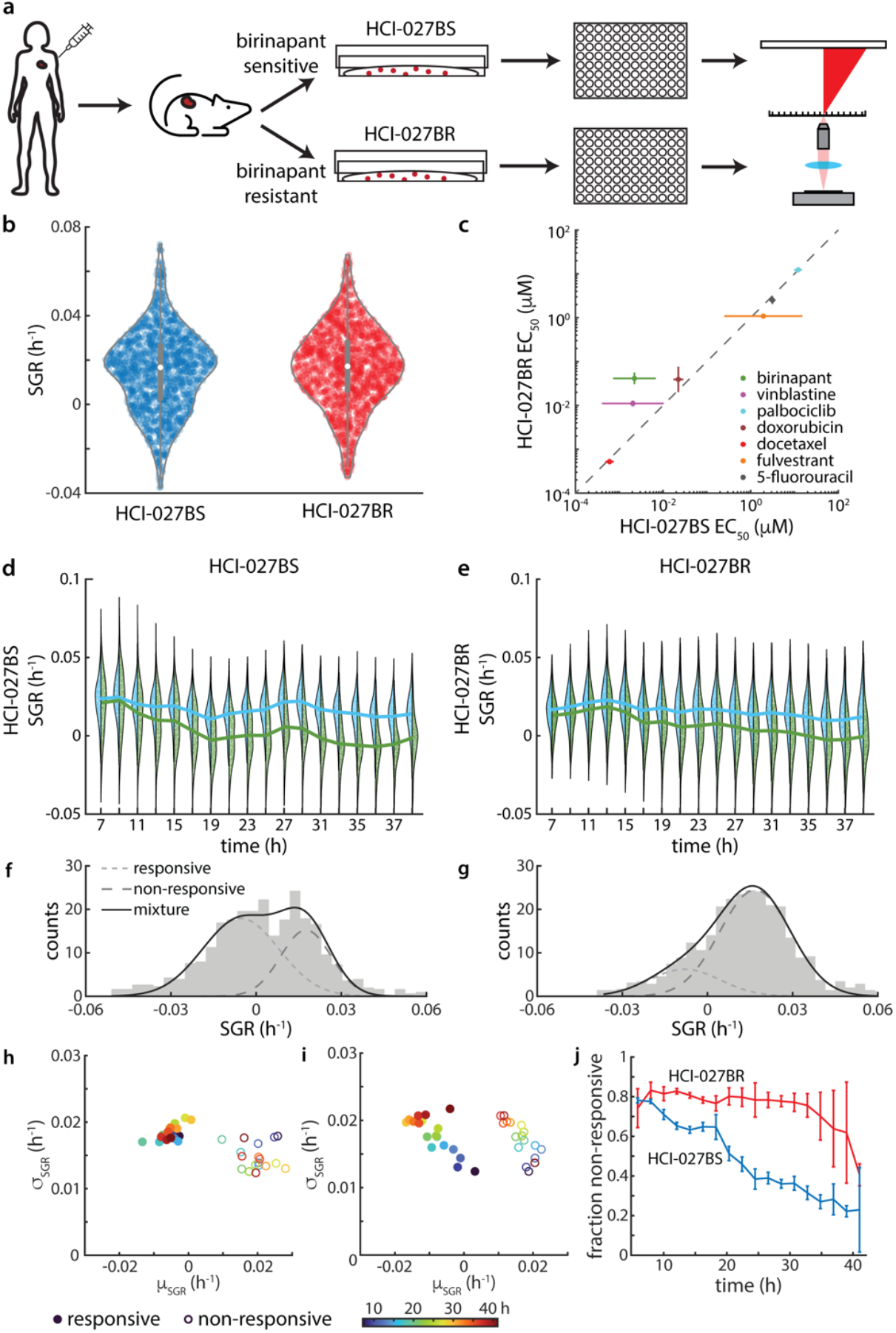
mQPI captures non-responsive subpopulation response in a heterogeneous sample. (a) Experiment workflow illustrating how models were derived and plated for QPI. (b) Violin plots of SGR for DMSO control, *p* = 0.014 from a paired sample t-test. (c) Comparison of EC_50_ for HCI-027BR and HCI-027BS. (d, e) Side by side violin plots showing population SGR over time for (d) HCI-027BS and (e) HCI-027BR. At each timepoint, the left side distribution (blue) shows untreated cells and the right side distribution (green) shows cells treated with 1.6 nM birinapant. (f,g) Subpopulations distributions of SGR for (f) HCI-027BS and (g) HCI-027BR treated with 1.6 nM birinapant at 23 h. Subpopulations estimated from Gaussian mixture modeling shown as solid lines (small dash: responsive, large dash: non-responsive, black: mixture, R^2^ = 0.98 panel f, R^2^ = 0.99 panel g). (h,i) Time evolution of mean (*μ*) and standard deviation (*σ*) of responsive (closed symbols) subpopulations and non-responsive (open symbols) subpopulations from Gaussian mixture modeling in (h) HCI-027BS and (i) HCI-027BR. Symbols are colored by time (blue to red). (j) Non-responsive fraction of the population from the fitted model.

EC_50_ measurements from the two models are strongly correlated to each other for most drugs (R^2^ = 0.916, *p* = 7×10^−4^) (Fig. 3c, S8, S9). However, as expected, HCI-027BS has a significantly lower EC_50_ than HCI-027BR in response to birinapant (EC_50,HCI-027BS_ = 2.2 nM, EC_50,HCI-027BR_ = 41 nM). To evaluate if the difference in response to birinapant is due to the presence of a resistant subpopulation, we examined the distributions of single cell responses to birinapant over time (Fig. 3d,e). These data show significant heterogeneity in birinapant treated populations (Fig. S10), especially at intermediate timepoints, for example at 23 h (Fig. 3f,g). We fit these single cell SGR distributions of birinapant treated cells using a Gaussian mixture model. This model includes two populations, each modeled with a Gaussian distribution of SGR: a higher growth rate corresponding to a non-responsive population and lower growth rate corresponding to a responsive population. Surprisingly, in response to 1.6 nM birinapant (just below the EC50 for HCI-027BS), both HCI-027BS and HCI-027BR show subpopulations of responsive and non-responsive cells over time (Fig. 3 h,i). In HCI-027BS, both the responsive and non-responsive subpopulations show no clear trend in growth rate or distribution width (Fig 3h). Additionally, the fraction of non-responsive cells in HCI-027BS decreases steadily over time (Fig. 3j). This suggests that cells in HCI-027BS show a delayed response, with no persistent, resistant subpopulation.

In contrast, in HCI-027BR, both responsive and non-responsive subpopulations show a trend towards decreasing growth rate, suggesting an impact in growth rate on both groups. However, the non-responsive subpopulation persists over the 45h experiment (Fig. 3j). This suggests the presence of a dominant and persistent resistant population in HCI-027BR, as expected based on its derivation from a birinapant resistant PDX model. Overall, these data demonstrate the ability of QPI to identify drug resistant subpopulations from mixed samples.

### Rapid assessment of response via mQPI on high purity, direct-from-thaw patient samples

Two pleural effusion samples, procured as fresh frozen and banked at Huntsman Cancer Institute, were tested by mQPI. Samples were thawed from liquid nitrogen storage and depleted of CD45+ cells via magnetic bead sorting to enrich for non-immune cells including tumor cells. We then plated the primary cells in 2D monolayer cultured in Clevers’ complete medium^60^ in a 96 well plate for 18 hours prior to drug treatments^9,59,61^ (Fig. 4a). Both samples were characterized via Hematoxylin and Eosin (H&E) staining (Fig. 4b) and immunohistochemistry (Figs. S11, S12) to confirm the presence of cells with tumor-like morphology and staining (Fig. 4b). When imaged using QPI the sample from patient 1 displayed two distinct cell sizes (Fig. 4c) and the overall mass distribution of single cells displays two distinct peaks (Fig. 4d). We used that distinction in mass apparent by the bimodal distribution in patient 1 (Fig. 4d) to filter out the smaller cells that did not grow in control conditions and likely represented residual immune and/or other non-tumor cells (Fig. 4e). This resulted in a clearly discernable drug response among the larger mass cells (Fig. 4f).

**Figure 4.**
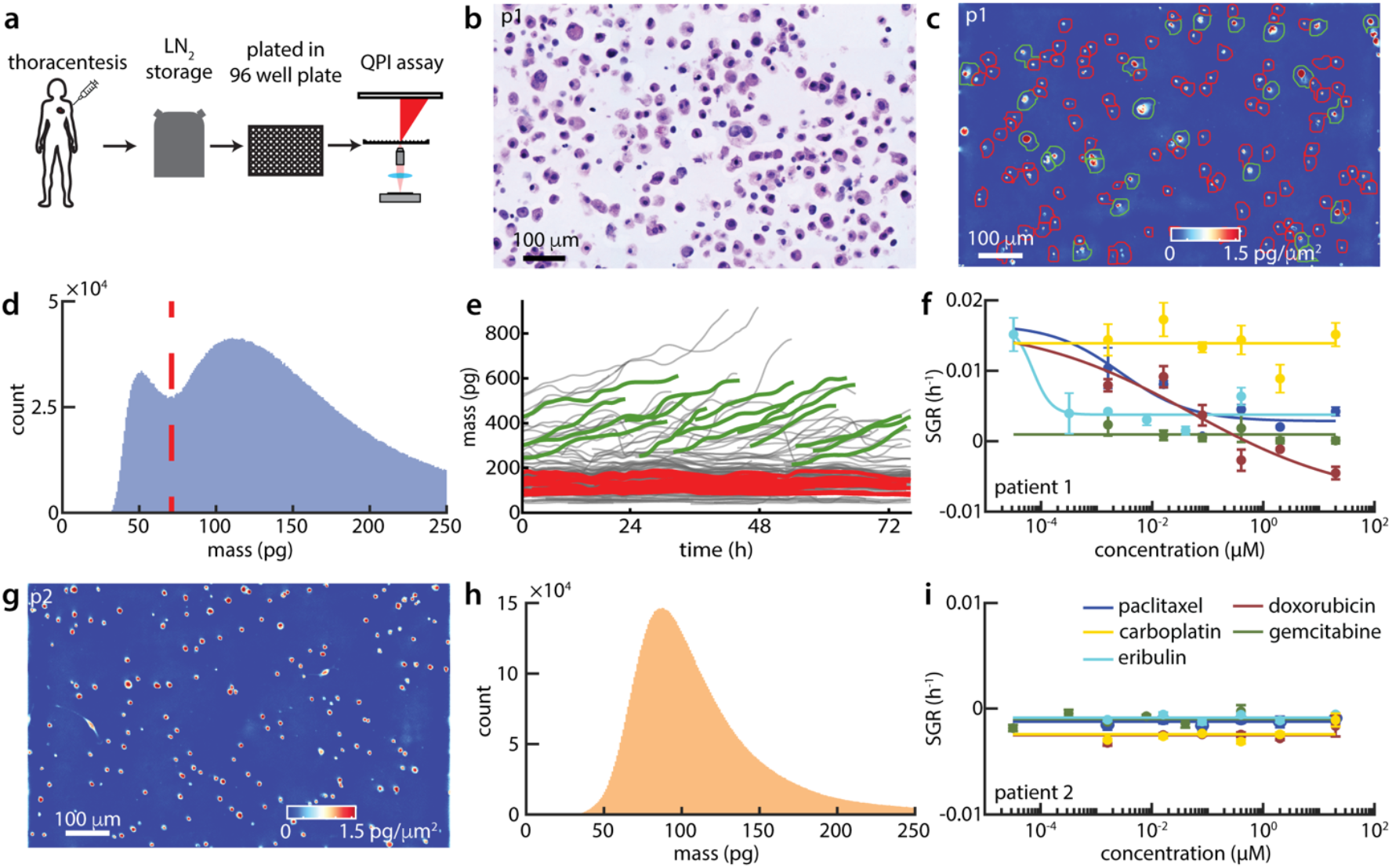
Direct-from-thaw patient samples analyzed using mQPI. (a) Patient sample workflow for QPI on patient pleural effusion samples direct-from-thaw after storage in liquid nitrogen (LN_2_). (b) H&E stain image for pleural effusion sample from patient 1 (p1). (c) QPI image for patient 1 cells from small cell enriched region. Green/red segmentation outlines indicate cells greater/less than the mass threshold shown in panel d. (d) Distribution of cell mass across all positions for patient 1. Red dashed line is a mass threshold. (e) Single cell mass over time for one well of vehicle (DMSO) control from patient 1. Select cells above/below the mass threshold are highlighted in green/red, respectively. (f) Drug response for patient 1, direct-from-thaw. (g) Representative QPI image of patient 2 cells. (h) Distribution of cell mass across all positions for patient 2. (i) Drug response for patient 2, direct from thaw.

The sample from patient 2, however, had cells that were more uniform in size, as observed in QPI images (Fig. 4g) and single cell mass distributions (Fig. 4h). This sample also had notably higher positive staining for CD45 (Fig. S12). Furthermore, we were unable to identify a subpopulation of cells with a differential response in the patient 2 direct-from-thaw sample, with all dosed conditions (Fig. 4i) and untreated control (SGR_patient2,DMSO_ = −2.2×10^−3^ h^−1^) both having a slightly negative SGR.

### Short expansion of primary cells enriches tumor population enabling analysis by mQPI

To allow cells to acclimate after thawing and to enrich the subpopulation of growing cells (Fig. 4e), we tested the effect of a short-term expansion by culturing primary cells in 2D with regular media changes for two weeks (Fig. 5a). This successfully increased the fraction of large, growing cells in both patient pleural effusion samples (Fig. 5b,c). Patient 1 drug response were consistent with results in the direct-from-thaw experiment (compare Fig. 4f and Fig. 5d), and visible in normalized mass over time as well (Fig. 5e). Similar to the maintenance of drug sensitivities observed for expansion of PDO in Matrigel,^59^ these responses were also maintained in 2D culture on PEI and Matrigel for at least 6 weeks (Fig. S14, S15). For patient 2, we were able to gauge a response after short-term 2D expansion that we were unable to observe in the direct-from-thaw setting, including doxorubicin sensitivity and mild stimulatory effects of paclitaxel (Fig. 5f). Overall, this proof of concept suggests that mQPI analysis can be applied to a broader range of patient samples with a short-term expansion period.

**Figure 5.**
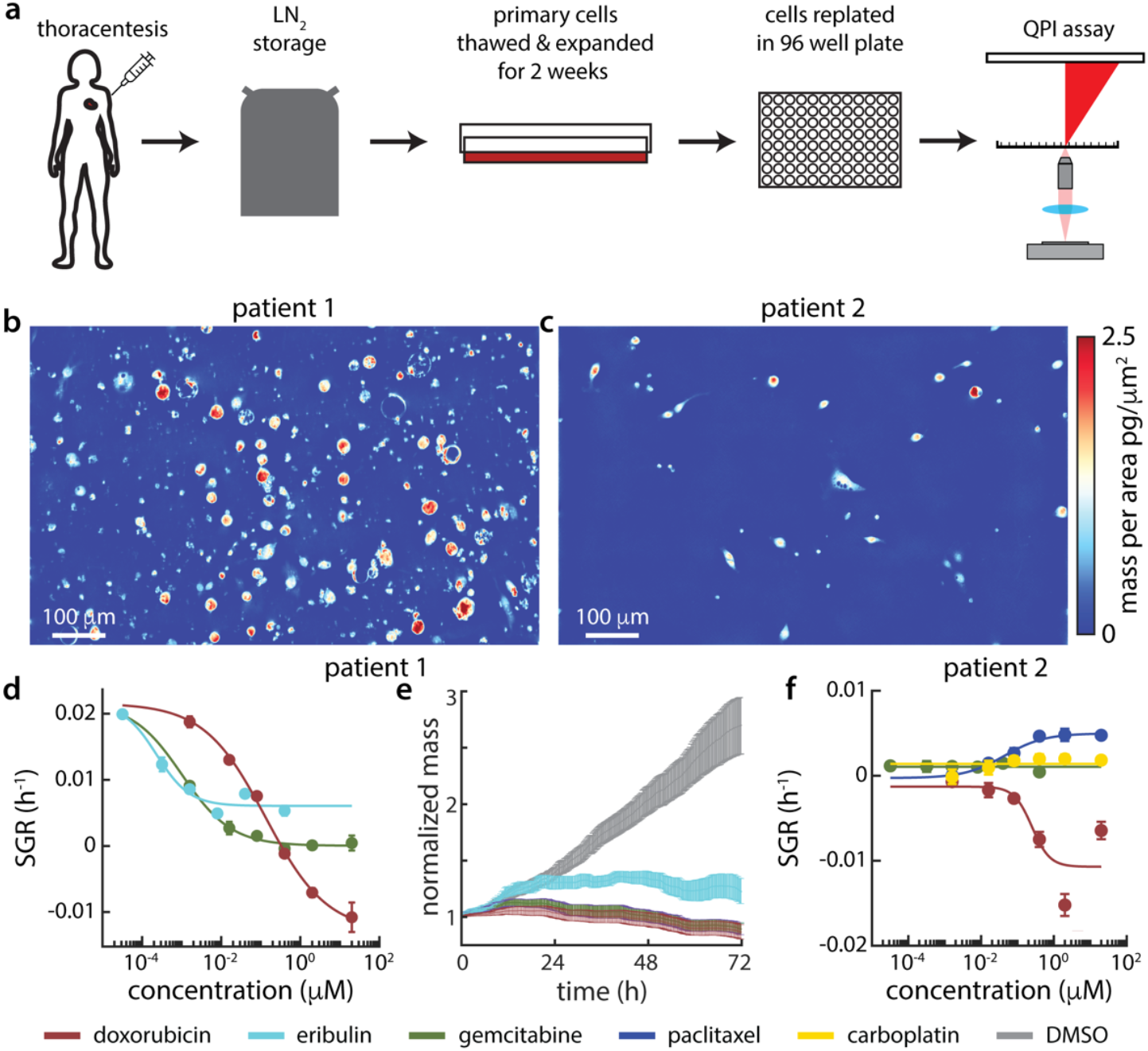
Short expansion enriches cell population for analysis by mQPI. (a) Workflow for pleural effusion patient sample expansion prior to QPI imaging. (b) QPI image of patient 1 cells after expansion. (c) Phase image for patient 2 cells after expansion. (d) Dose response for patient 1 after expansion. (e) Normalized mass over time for patient 1 cells after expansion. (f) Dose response for patient 2 after expansion. Colors corresponding to drugs for (e-f) indicated at the bottom of the figure.

## Discussion

We demonstrated that mQPI can provide functional cell data for PDMCs, including both PDXO cells and cells from viably frozen patient samples. For PDXOs originating from the same patient, mQPI could identify different drug sensitivity profiles based on anatomic site and/or time of collection (Fig. 2), which underscores the importance of accounting for site-specific tumor heterogeneity when selecting therapies. The ability of mQPI to track single cells also allows for heterogeneity to be directly observed. In application to a sample with mixed responses (Fig. 3), mQPI was able to distinguish resistant subpopulations, in contrast to conventional, bulk assays, which would be unable to resolve single cell responses. Additionally, as in previous work,^53^ mQPI measurements (ToR, DoR, EC50, SD) had low correlation to one another, indicating that we were able to capture independent measurements of response to therapy in PDMCs (Fig. S15).

We also demonstrated the ability of mQPI to detect response from viably frozen patient samples, which could represent a key step toward practical application of mQPI in the clinical setting. Our overall assay time was either days (direct-from-thaw, Fig. 4) or 2.5 weeks (Fig. 5), which is a significant improvement over the months often required to establish PDX or PDO models. We found that patients’ samples tend to be contaminated with different cell types even after CD45 depletion, but were able to use a mass-based filter to computationally increase the percentage of tumor cells evaluated and gauge a tumor cell response in mixed cultures. However, this filtering approach alone was not sufficient for both samples we tested. Future work will examine alternative filtering methods including label-free and machine learning/AI-assisted approaches based on cell morphology or other features accessible via QPI. Our data indicate mQPI is able to capture a response from a very small subset of cells, addressing a key limitation of functional assays in clinical settings. It is worth noting that for this study, we plated approximately 500,000 cells per experimental condition, but performed imaging on approximately 50,000 cells. This represents just a 10% sample utilization, which can likely be further improved with optimized sample handling.

In summary, mQPI is a rapid, accessible, and robust tool for analyzing cancer cell responses to therapies, offering insights into growth, heterogeneity, and drug sensitivity. To enhance clinical relevance, future efforts should focus on optimizing tumor cell enrichment, whether through improvements in sample preparation or through computational filtering, and should be extended to comparison with known clinical outcomes in larger sample cohorts. The ability to rapidly assess therapeutic responses using direct-from-thaw samples or after short-term expansions underscores its potential as a time-efficient alternative to current models. Ultimately, mQPI holds promise for advancing functional precision oncology and improving personalized treatment strategies.

## Methods

### 96-well plate preparation

10x plate coating solution was prepared by diluting 25,000 MW branched polyethylenimine (PEI, Millipore Sigma, USA) at 0.1% in borate buffer (pH 8.4). Immediately prior to plating, 10x PEI was then diluted in borate buffer to reach 0.01% working concentration. 96-well plates (Eppendorf, Germany) were treated with 700 μL of 0.01% PEI per well for 4-12 h then washed twice with autoclaved deionized water. Plates were either used immediately or stored up to two weeks at 4°C.

### PDXO model culture

The PDXO models were established in the Welm labs at the Huntsman Cancer Institute at the University of Utah and were cultured as described previously^59^ for 1-2 passages (2-8 weeks) post thaw prior to plating for QPI. 18-24 hours prior to imaging via QPI, PDXOs were dissociated into single cell suspension as described previously^59^ and plated at a density of 5,000 cells per well using PDXO media^59^. The moat in the 96-well plate was filled with sterile, deionized water to minimize evaporation (8 mL inner moat and 5 mL outer moat).

### Primary cell culture

Patient samples were thawed, washed and resuspended at a maximum of 1 × 10^8^ cells/mL in cold PBS containing 2% fetal bovine serum (FBS) and 1 mM EDTA. Cells were immune cell depleted by anti-using EasySep CD45 antibody Depletion Kit II (StemCell, Cat # 17898) (1x for patient 1, 2x for patient 2). Cells were then pelleted and resuspended in Clevers media^60^ then plated at 12,000-15,000 cells per well in PEI coated 96-well plates, or plated for 2-week expansion with one intermediate media change, then plated for QPI. Fresh media was provided immediately prior to treatment and imaging by QPI.

### Drug dosing

Cells were dosed (Table ST2, Selleck Chemicals, USA) 3 h prior to the start of imaging by adding 100 μL at 2x the dosing concentration diluted in media to wells containing 100 μL of media. Vehicle (DMSO or ethanol) concentrations were maintained at 0.1% across all wells to account for potential vehicle toxicity.

### Quantitative Phase Imaging

QPI was performed using a custom DPC QPI microscope described and characterized previously.^53^ The microscope is optimized to rapidly acquire data from 9 locations in each well of a 96-well plate (864 total locations) every 20 minutes. Experiments were performed in triplicate. The microscope uses an LED array for illumination (Adafruit, New York, USA) to rapidly switch between half circle illumination,^34,35,62^ a 10x Olympus Plan N objective lens (Olympus, Japan) with a numerical aperture of 0.25, a tube lens, and a Grasshopper3 machine vision monochromatic camera (Teledyne-FLIR, Oregon, USA) with an exposure time of 50 ms and gain of 25 dB. An MLS203 high-speed stage (Thorlabs, New Jersey, USA) is used for stage motion. The microscope is compact enough to operate inside of a tissue culture incubator to maintain cell culture conditions at 37 °C, 5% CO_2_, and 95% humidity. Raw DPC intensity images were processed for quantitative phase using Tikhonov regularization^35,62^ with a regularization parameter of 1×10^−3^.

### Image processing

Image processing was completed using methods and code as described previously^30^ (https://github.com/Zangle-Lab/MultiparametricQPI). Briefly, cells were segmented by Sobel filtering to find edges, followed by morphological processing to dilate segmented cell areas. Cell mass was computed using a specific refractive increment of 1.8×10^−4^ m^3^/kg^63^. Cell tracking was performed by minimizing distance between paired cells^64^ in x,y, position as well as mass.

### Growth rate Processing

Specific growth rates (SGRs) were computed by least squares linear fitting to tracks longer than 20 frames after applying a median filter of size 5 frames. SGR was computed as the slope of this linear fit divided by the initial projected mass at the start time of the track. SGR data for each condition were averaged and used for Hill curve fitting:

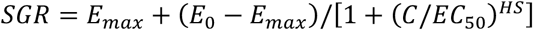

where *E*_*max*_ is the upper asymptote, *E*_*0*_ is the lower asymptote, *C* is drug concentration, *EC*_*50*_ is the inflection point of the Hill curve, and *HS* is the Hill slope. Depth of response (DoR) was then defined as:

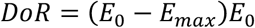

### Time of response calculation

To compute time of response (ToR), cell mass data were binned into 12 h windows centered on time points at 2 h increments. Slope was extracted from the model and normalized by the initial mass predicted by the linear fit, as in overall SGR calculations. Hellinger distance between two populations was then computed from the probability density functions of SGRs of those populations.^65^ The Hellinger distance threshold for response was defined as the maximum Hellinger distance between the two on-plate controls for each experiment. For each drug treatment, we measured the Hellinger distance at each time point between the drug treated distribution and the control distribution and fit an exponential model to these data. ToR is defined as the time when the curve fit to the measured Hellinger distance crosses the threshold determined from control data.

### Gaussian mixture analysis

Mean SGR values at each time point were binned into probability histograms and smoothed using kernel density estimation (KDE) with a bandwidth of 0.0025 h^−1^. KDE curves were then fit to a sum of two gaussians, or gaussian mixture model:

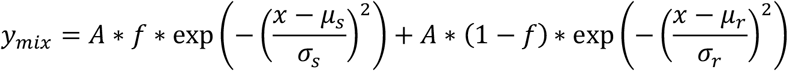

with *f* representing the fraction of the overall population that is sensitive and *A* the amplitude of the Gaussian distribution. *μ* and *σ* indicate the mean and standard deviation of the two Gaussian distributions, where subscripts indicate responsive, *r*, or non-responsive, *n*, populations. Initial fitting parameters for the first time point were estimated from the KDE amplitude (*A*_*KDE*_) the median (*m*_*all*_) and the standard deviation (*σ*_*all*_) of the overall population: *A*_*0*_= *A*_*KDE*_; *μ*_*r,0*_ = *m*_*all*_ -*σ*_*all*_/2; μ_*n,0*_ = *m*_*all*_ + *σ*_*all*_/2; *σ*_*t,0*_ = *σ*_*n,0*_ = *σ*_*all*_. The resulting parameters from each fit were then used as initial guesses for the next time point.

### Statistics

Group means were compared using a two-tailed Student’s t-test as implemented in MATLAB’s ttest2 function (MathWorks, Natick, MA, USA). Welch’s correction for unequal variances was applied. Variances between fits were compared using MATLAB’s ftest function, which performs a two-tailed F-test. Significance was set at *p* < 0.05. Lin’s concordance coefficient was used to measure the concordance between variables.^66^

## Acknowledgements

This work was conducted with funding from the Department of Defense Breast Cancer Research Program (W81XWH1910065; T.A.Z./P.J.B., HT94252410020; T.A.Z., and W81XWH1210077; A.L.W.); the National Cancer Institute (U54CA224076; A.L.W. and U01CA217617; B.E.W.); and the Breast Cancer Research Foundation Founders Fund (A.L.W.). Research reported in this publication utilized the Biorepository and Molecular Pathology Shared Resource and the Immuno-Oncology Network initative at Huntsman Cancer Institute at the University of Utah, supported by the National Cancer Institute of the National Institutes of Health under Award Number P30CA042014. The content is solely the responsibility of the authors and does not necessarily represent the official views of the NIH.

