## Supporting Information for "Determination of Drug Sensitivity in Patient Derived Models of Breast Cancer by Multiparametric QPI"

**Contents:**

Supplemental Tables ST1-ST3
Supplemental Figures S1-S15

**Table ST1. Models used for this study and their receptor status.**

| Model Name | ER | PR | HER2 |
|---|---|---|---|
| **HCI-27BS/BR** | **-** | **-** | **-** |
| **HCI-37** | **-** | **-** | **-** |
| **HCI-38** | **-** | **-** | **-** |
| **HCI-39** | **-** | **-** | **-** |
| **Patient 1** | + | **-** | **-** |
| **Patient 2** | + | **-** | **-** |

**Table ST2. Treatments used in this study, their targets, abbreviations, and dose range.**

| Drug Name | Abbreviation | Target | Dose Range |
|---|---|---|---|
| **Birinapant** | biri | cIAP1 and cIAP2 | Low* |
| **Vinblastine** | vin | Microtubule | Low* |
| **Palbociclib** | palbo | CDK4/6 | High |
| **Doxorubicin** | dox | DNA topoisomerase II | High |
| **4-Hydroxytamoxifen** | 4HT | Estrogen receptor | High |
| **Docetaxel** | doc | Microtubule | Low |
| **Lapatinib** | lap | EGRF/HER2 | High |
| **Fulvestrant** | fulv | Estrogen receptor | High |
| **Carboplatin** | carbo | DNA synthesis | High |
| **5-Fluorourcil** | 5FU | DNA-RNA synthesis | High |
| **Paclitaxel** | pacl | microtubules | Low |
| **Eribulin** | eri | *microtubule* | Low* |
| **Gemcitabine** | gem | DNA synthesis & replication | High |

**Table ST3. Treatments associated with different panels and their corresponding number on the plate.** High range: 20, 2, 0.4, 0.08, 0.016, 0.0016 μM, Low range: 0.4, 0.04, 0.008, 0.0016, $3.2x10^{-4}$, $3.2x10^{-5}$ μM * initial testing performed in high range then switched to low to resolve response

| Panel | Panel A | Panel B | Panel C | Panel D |
|---|---|---|---|---|
| **Control 1** | Ethanol | DMSO | Untreated | Untreated |
| **Control 2** | DMSO | Untreated | DMSO | DMSO |
| **Drug 1** | Birinapant* | Docetaxel | Paclitaxel | Birinapant |
| **Drug 2** | Vinblastine* | Lapatinib | Gemcitabine | Gemcitabine |
| **Drug 3** | Palbociclib | Fulvestrant | Eribulin* | Eribulin |
| **Drug 4** | Doxorubicin | Carboplatin | Doxorubicin | doxorubicin |
| **Drug 5** | 4-Hydroxytamoxifen | 5-fluorourcil | Carboplatin | Carboplatin |

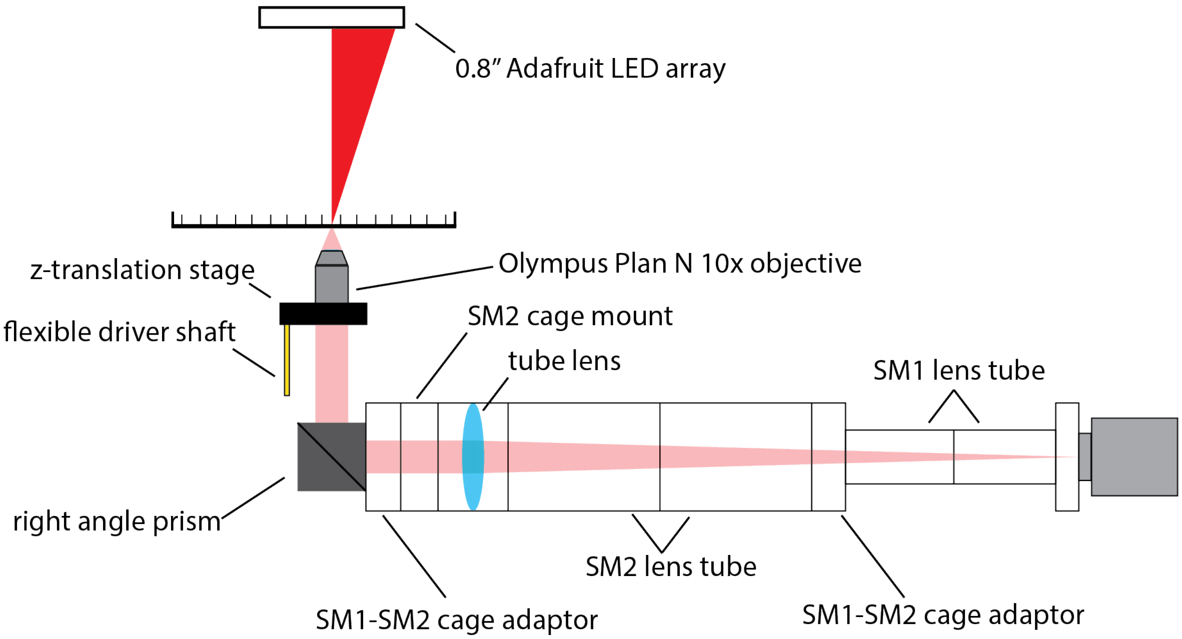


**Figure S1. Diagram of the custom QPI microscope used for imaging.**

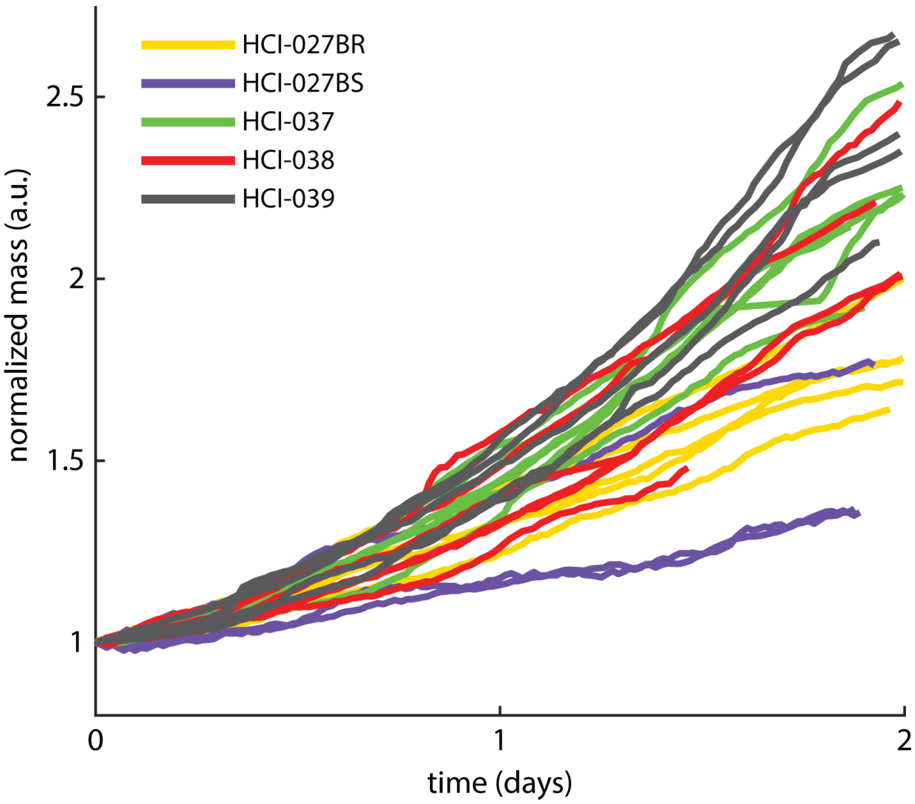


**Figure S2. Normalized mass of DMSO control for all tested PDxO models.**

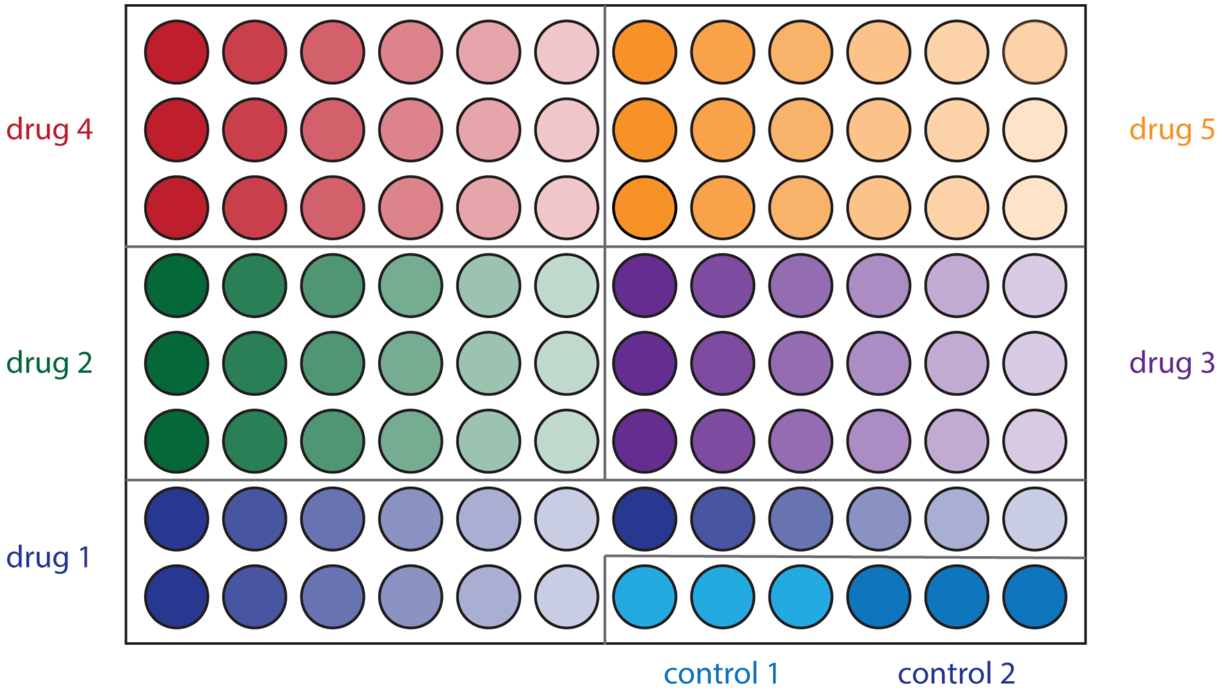


**Figure S3. 96-well plate layout.** Five treatments and arrangement of 6-point dose range for each is shown. Darker color (as on left side of each dose range) indicates higher concentration.

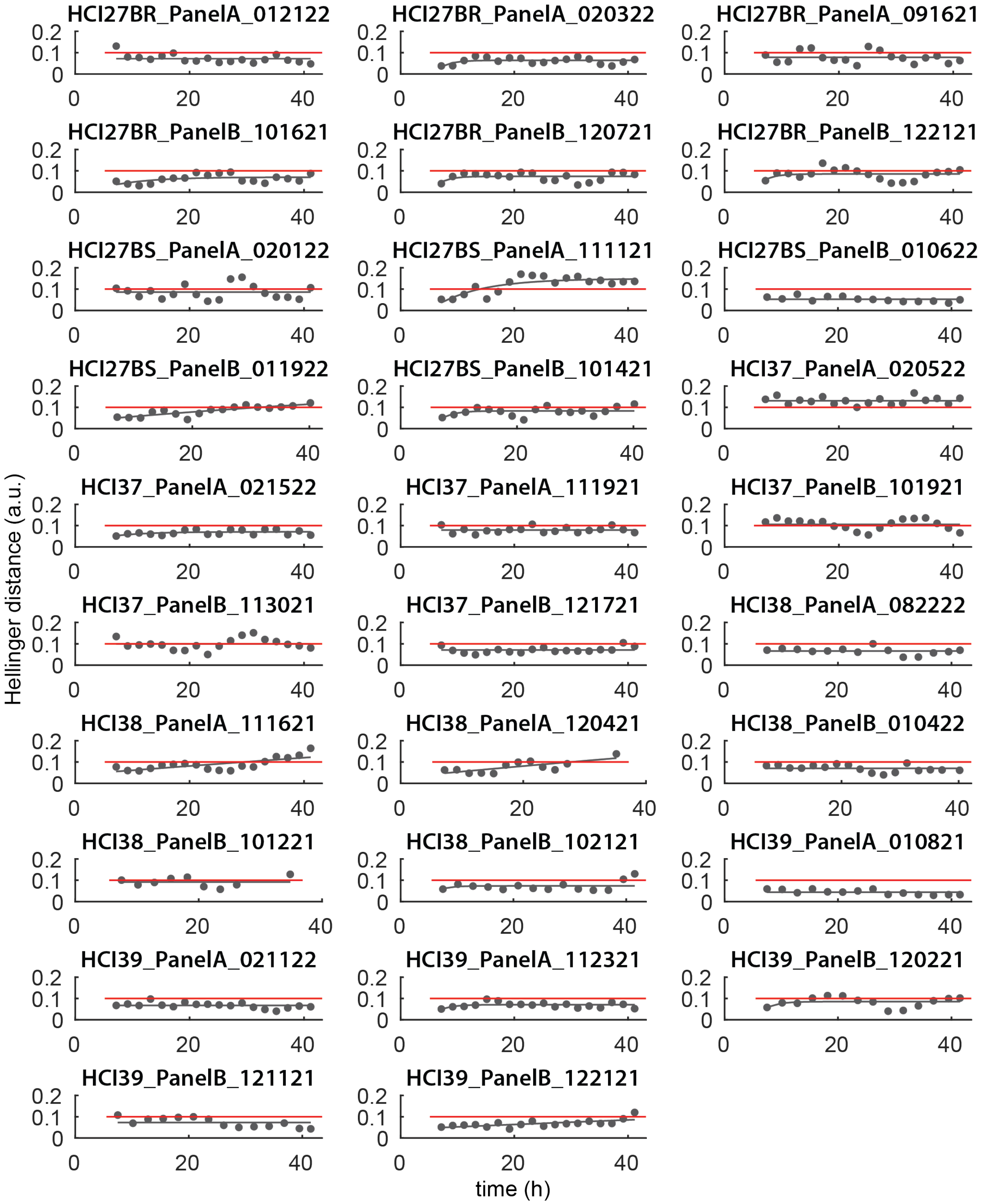


**Figure S4. Hellinger distance of DMSO treated control vs time.** The red line is the threshold at 0.1. Scatter dots are experimental data, and solid gray line is an exponential fit.

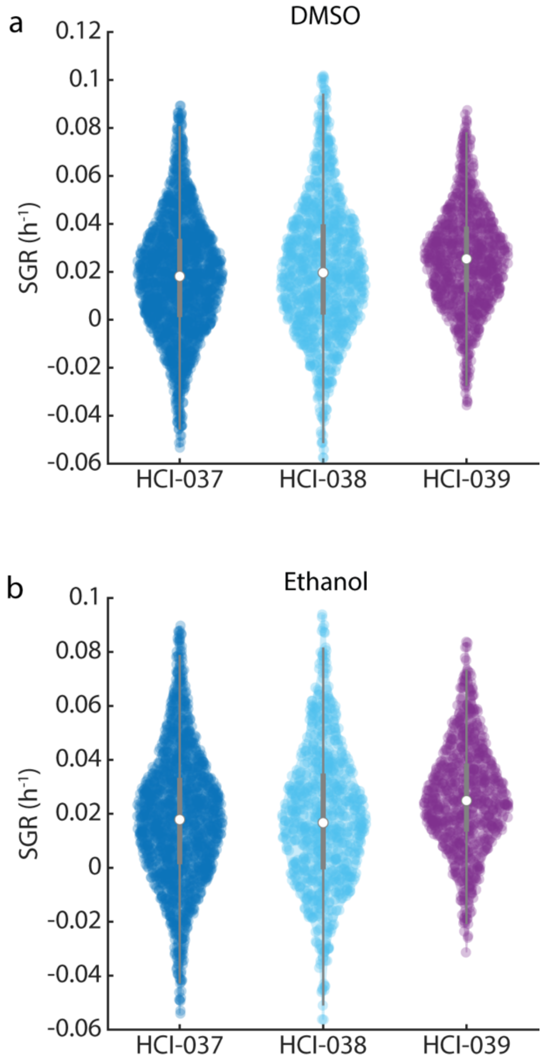


**Figure S5. Single cell growth rate distributions for HCI-037, HCI-038, and HCI-039 in vehicle control.** (a) DMSO, (b) Ethanol. Center point shows median. Grey bar indicates interquartile range.

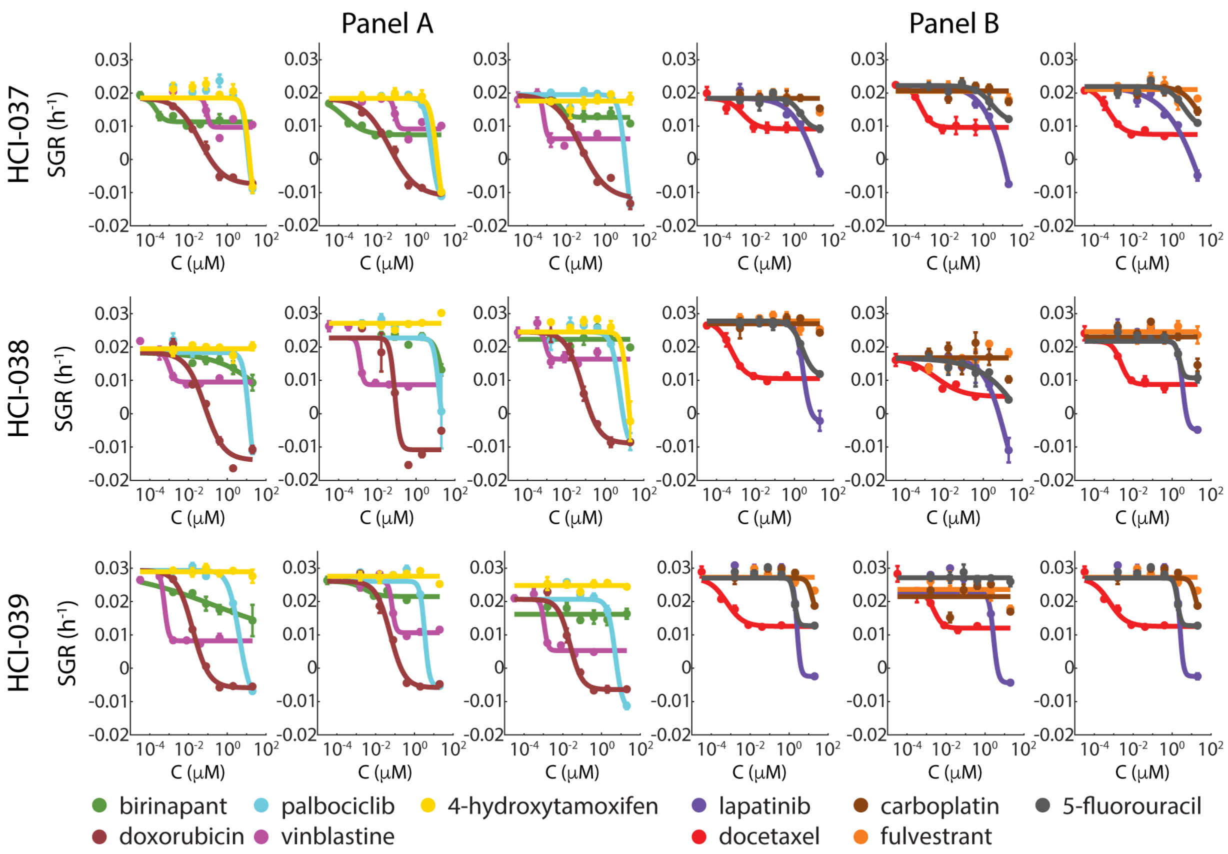


**Figure S6. Dose response plots for HCI-037, HCI-038, and HCI-039**. All replicates shown. Error bars represent SEM of individual wells.

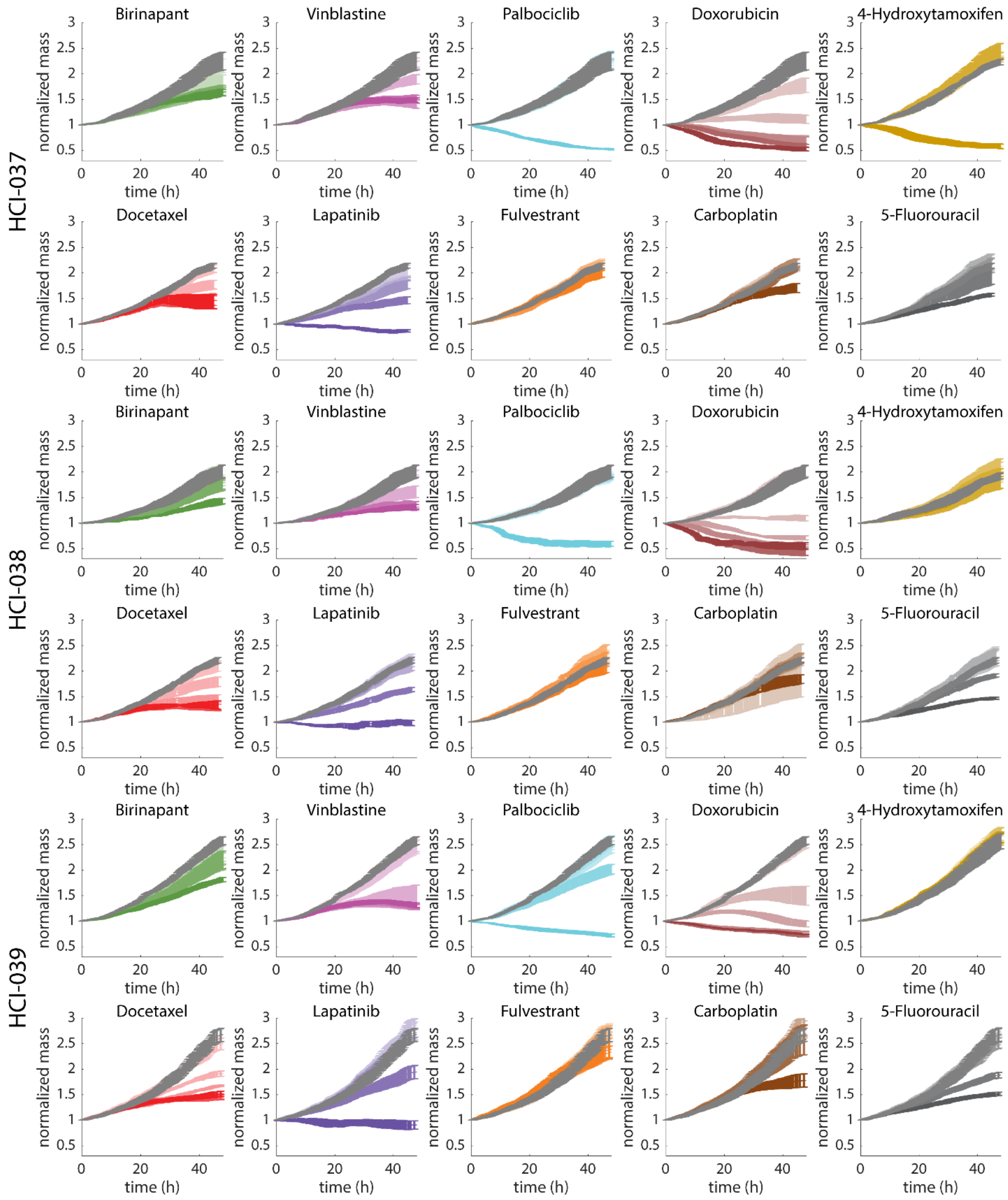


**Figure S7. Normalized mass vs. time data for HCI-37, HCI-38, and HCI-39.** Shading indicates concentration over the ranges given in ST2 and ST3. Darker color is higher concentration. Controls in gray. One representative replicate shown for each model. Error bars show SEM over individual imaging locations.

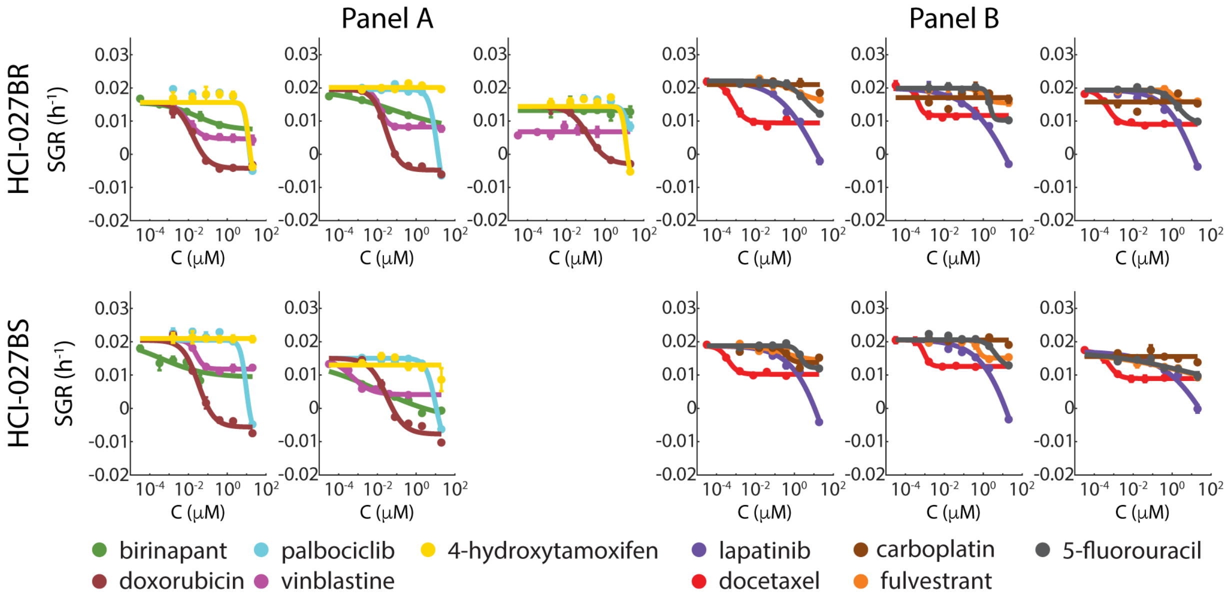


**Figure S8. Dose response plots for HCI-027BS and HCI-027BR.** All replicates shown. Error bars represent SEM of individual wells.

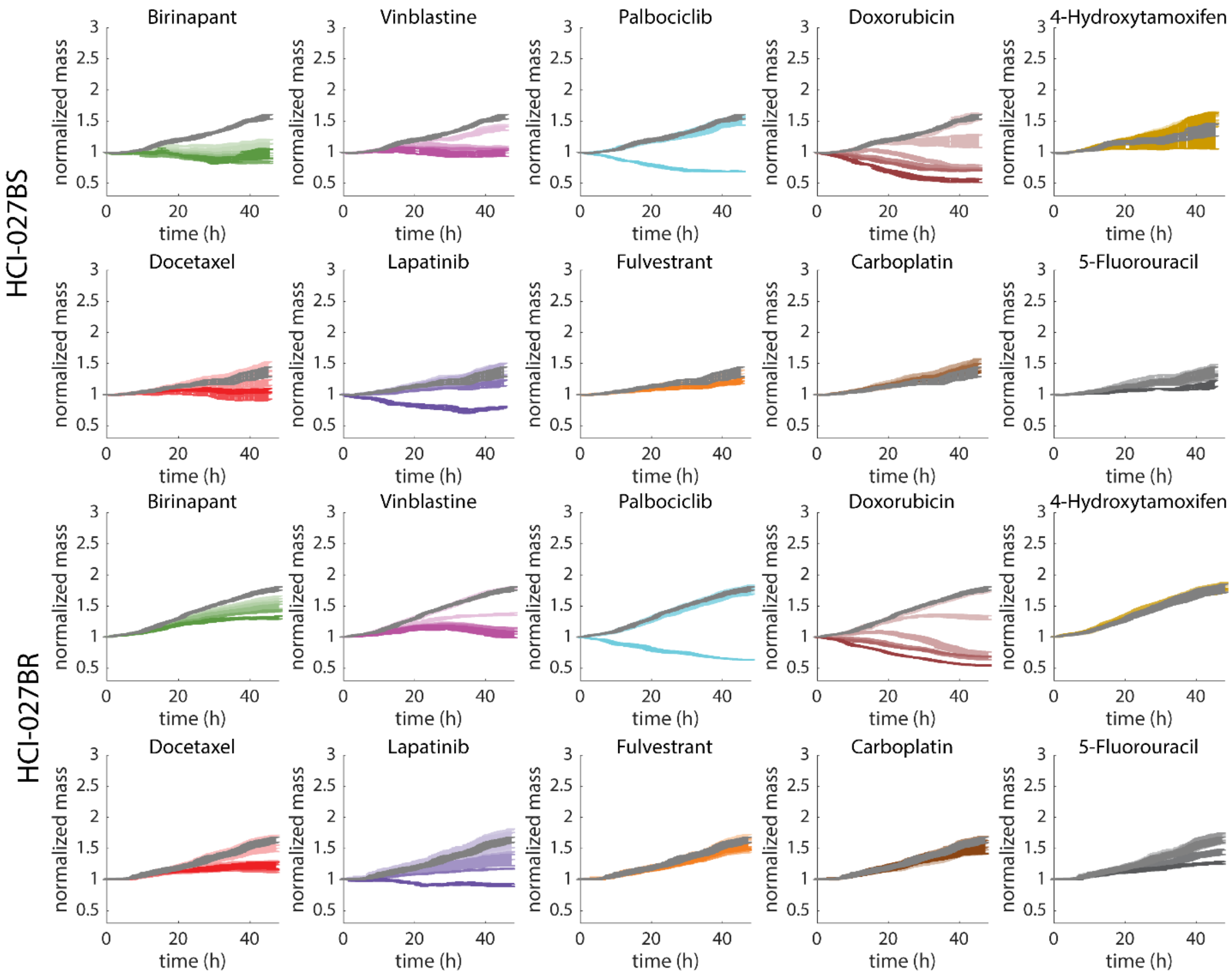


**Figure S9. Normalized mass plots for HCI-027BS and HCI-027BR.** Shading indicates concentration over the ranges given in ST2 and ST3. Darker color is higher concentration. One representative replicate shown for each model. Error bars show SEM over individual imaging locations.

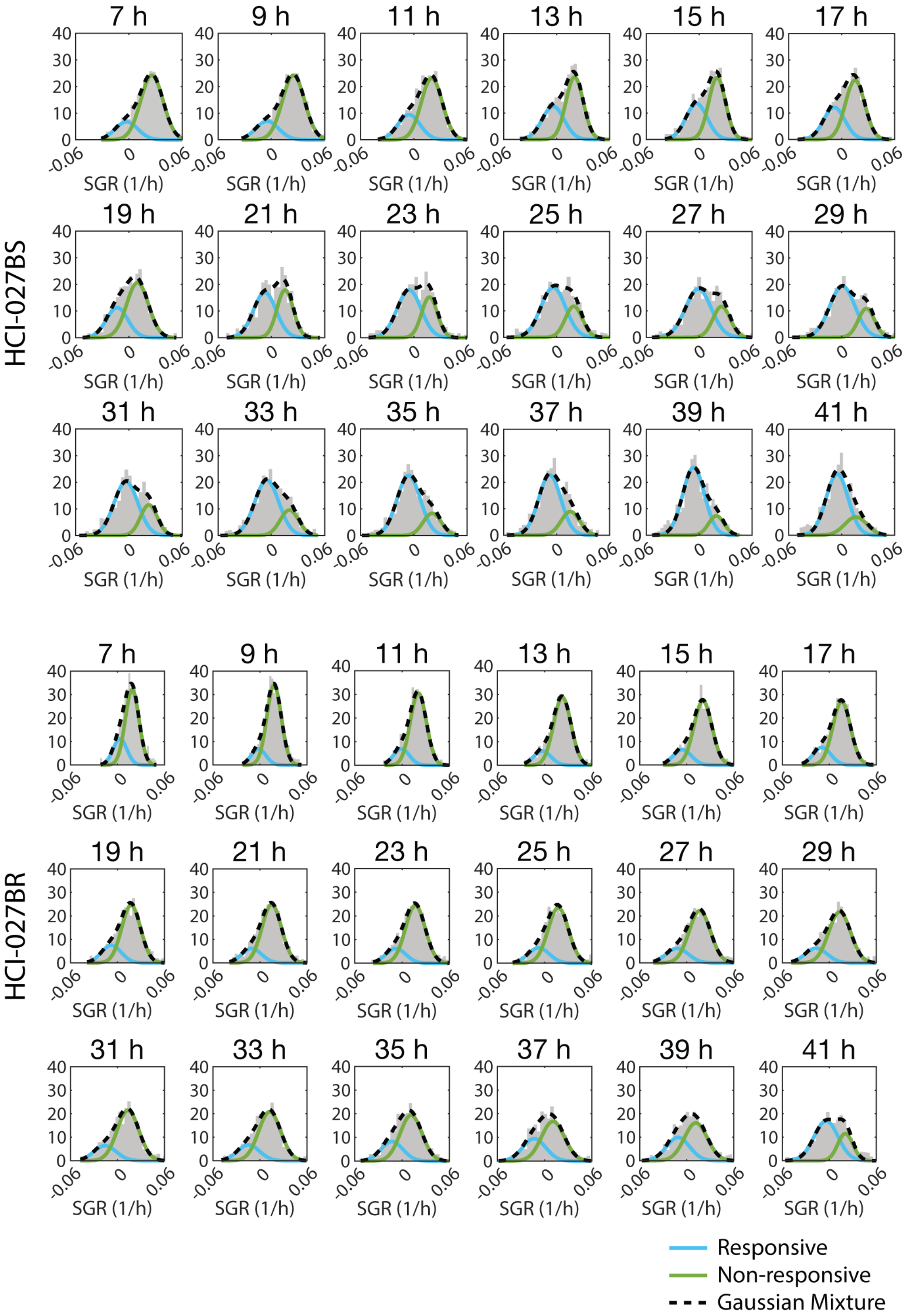


**Figure S10. Gaussian mixture model fits to HCI-027BS and HCI-027BR.** Histograms of specific growth rate (SGR) for individual cells at each timepoint shown in grey. Responsive and non-responsive population fits shown in blue and green, respectively. Black dashed line shows total model fit. All conditions treated with 1.6 nM birinapant.

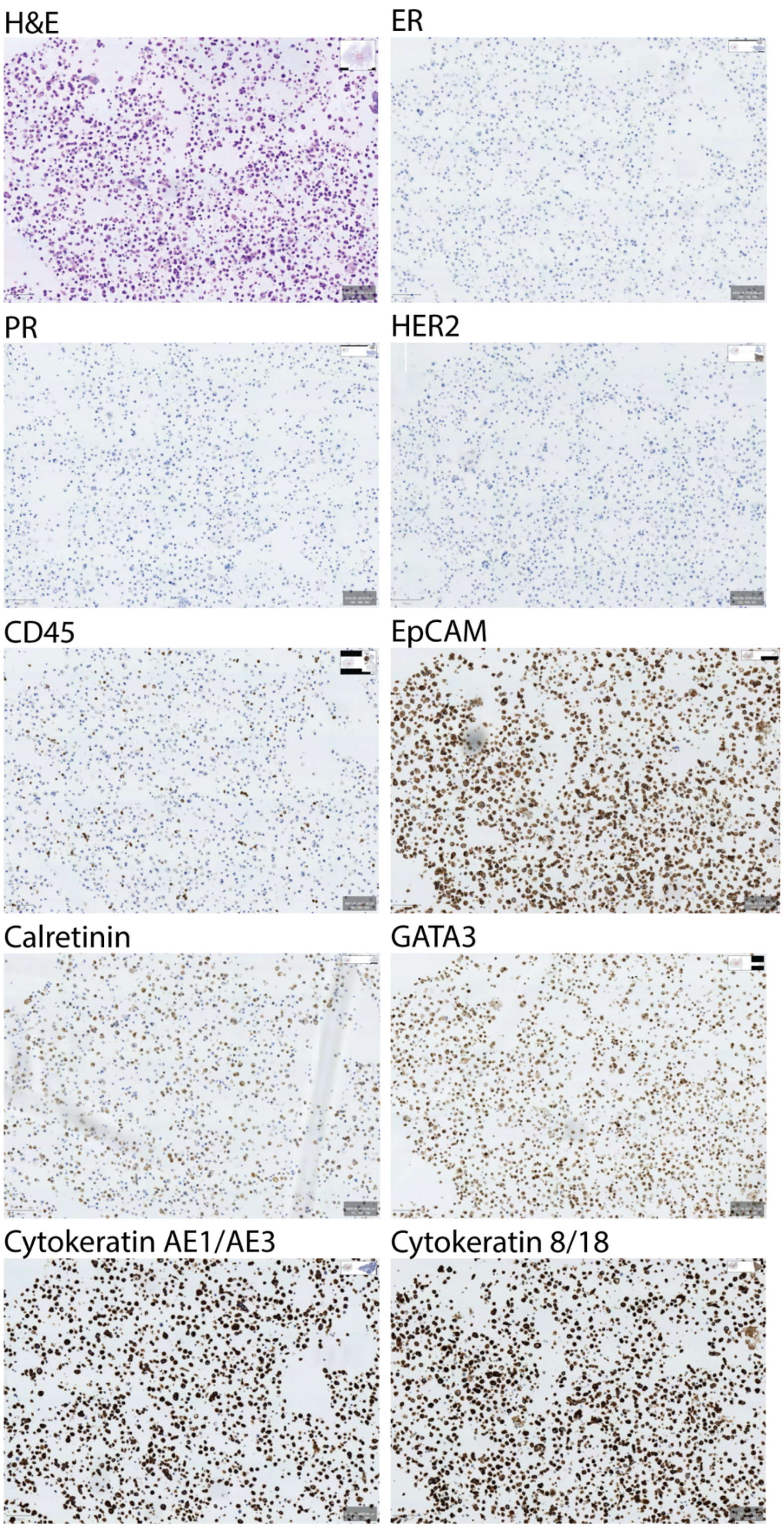


**Figure S11: Immunohistochemistry panel for patient 1.**

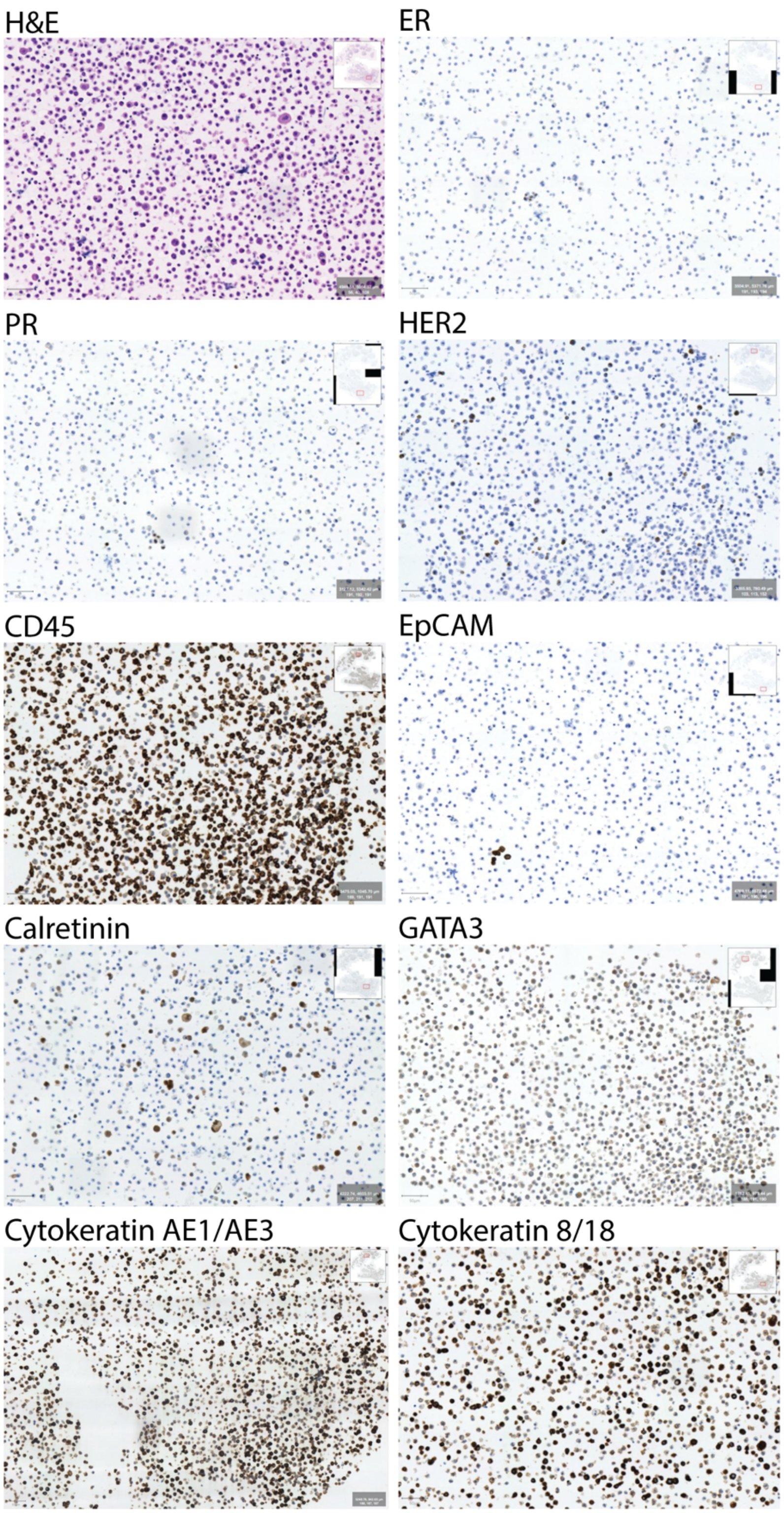


**Figure S12: Immunohistochemistry panel for patient 2.**

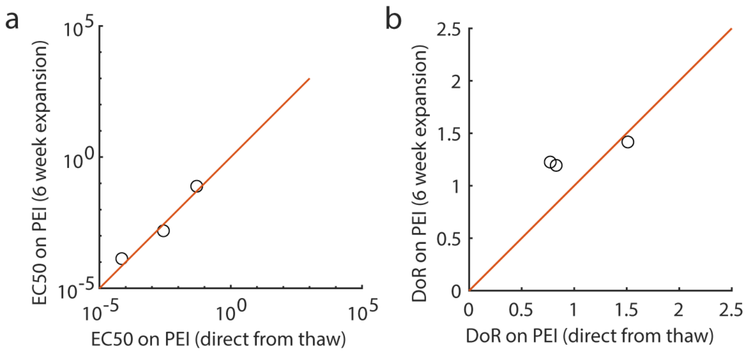


**Figure S13: Patient 1 responses direct from thaw compared to 6 week expansion**. (a) Comparison of EC50 values. (b) Comparison of depth of response (DoR)

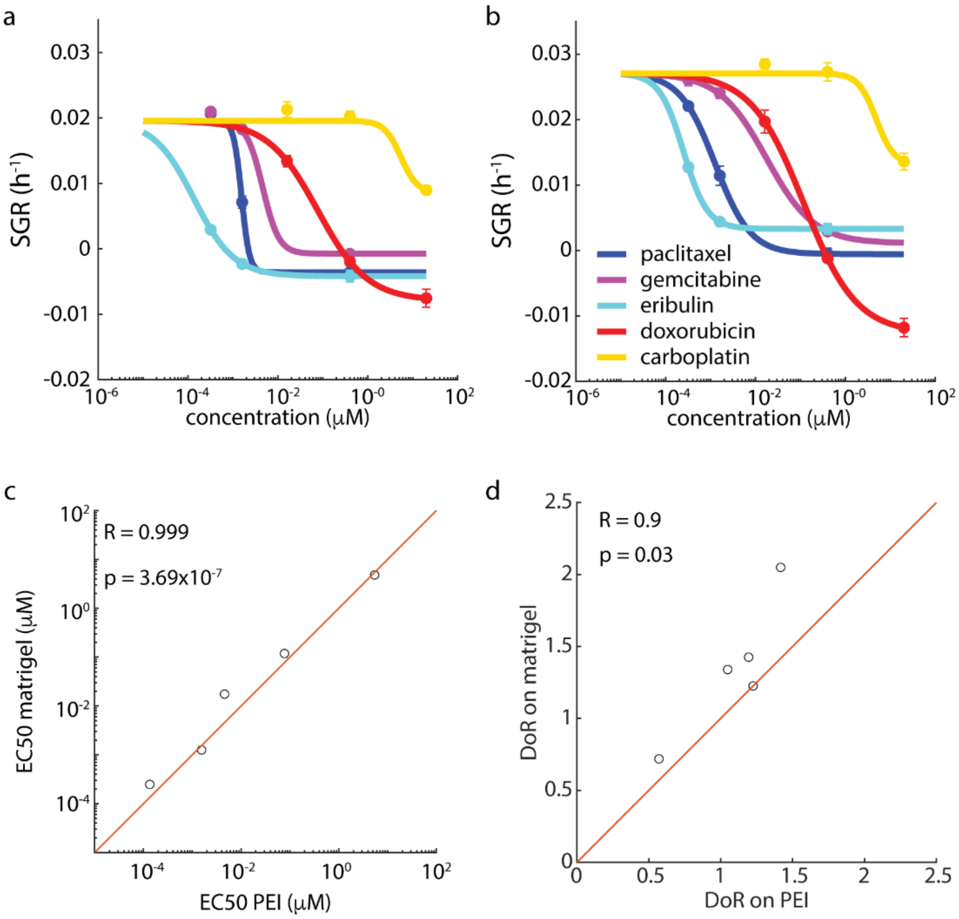


**Figure S14**. **Comparison of drug screening results on primary cells after 6 week expansion.** QPI dose response curves for cells expanded on (a) PEI and (b) Matrigel. (c) EC50 matches closely between the two condition (concordance coefficient = 0.99). (d) DoR for matrigel vs. PEI expansion (concordance coefficient = 0.70).

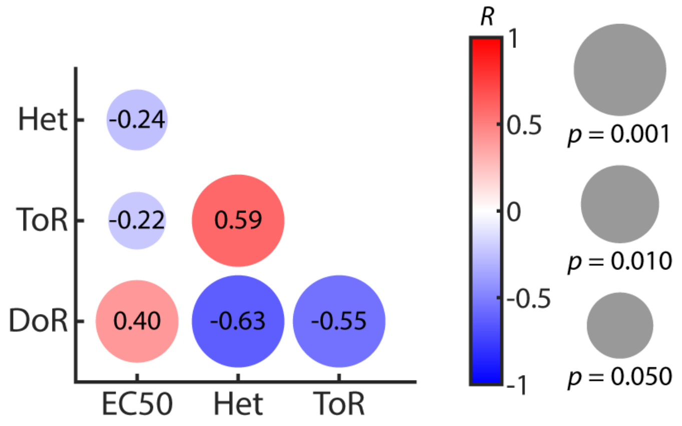


**Figure S15. Correlogram of mQPI parameters for tested PDMCs.** Color indicates Pearson's correlation coefficient, *R*. Size indicates *p*-value for the indicated correlation. Het = heterogeneity, ToR = time of response, DoR = depth of response.
